# Investigation of the transcriptional profile of human kidneys during machine perfusion reveals potential benefits of haemoadsorption

**DOI:** 10.1101/521294

**Authors:** John R. Ferdinand, Sarah A. Hosgood, Tom Moore, Christopher J. Ward, Tomas Castro-Dopico, Michael L. Nicholson, Menna R. Clatworthy

## Abstract

Transplantation is the optimal treatment for most patients with end stage kidney disease but organ shortage is a major challenge. Normothermic machine perfusion (NMP) has been used to re-condition marginal organs but the mechanisms by which NMP might benefit transplant kidneys are not fully understood. Furthermore, the question of whether removal of pro-inflammatory mediators from the perfusate might offer additional benefits in optimising kidneys prior to transplantation has not been addressed. Using pairs of human kidneys obtained from the same donor, we compared the effect of NMP with that of cold storage on the global transcriptome of kidneys, and then went on to investigate the impact of adding a haemoadsorption device to the NMP circuit. We found that cold storage significantly reduced the expression of inflammatory genes, but also of genes required for energy generation such as those encoding oxidative phosphorylation (OXPHOS) enzymes. In contrast, during NMP, there was marked upregulation OXPHOS genes, as well as a number of immune and inflammatory pathway genes. The induction of inflammatory genes during NMP was substantially attenuated by the addition of a haemoadsorber to the perfusion circuit, which also further increased OXPHOS pathway gene expression. Together, our data suggest that absorption of pro-inflammatory mediators from the perfusate represents a useful intervention that may further improve organ viability and should be tested in clinical practice.

**Single sentence summary:** The use of a haemoadsorber during machine perfusion reduces inflammatory gene expression, with potential benefits for kidney transplantation.

## Introduction

Kidney transplantation represents the optimal treatment for most patients with end stage kidney disease, with benefits for both quality and quantity of life (*1*). One of the current challenges in transplantation is organ shortage, leading to substantial waiting times for many patients. Several strategies have been employed to increase the number of kidneys available, including the use of deceased circulatory death (DCD) donors and extended criteria donors (ECD), both of which are associated with higher rates of delayed graft function (DGF) compared with deceased brainstem death (DBD) donor (*2, 3*). In DCD kidneys, exposure to warm ischaemia during the process of circulatory cessation makes a significant contribution to DGF. DGF occurs due to ischaemic tubular cell damage or death, which can stimulate innate immune activation via NLRP3 inflammasome assembly leading to the generation of interleukin(IL)1β and IL18, a process termed ‘sterile inflammation’ (*4–6*). Indeed, the presence of inflammatory cytokines in urine has been used as a biomarker of acute kidney injury and DGF (*7–9*).

Normothermic machine perfusion (NMP) allows transplanted organs to be perfused with warm, oxygenated red blood cells, in the absence of the immune components normally present in blood, including complement and neutrophils, with the aim of reversing the deleterious effects of warm and cold ischaemia (*10–14*). This process has been used to assess marginal organs (*15*) and to ‘re-condition’ organs to facilitate the transplantation of kidneys that were initially declined following offer via standard allocation schemes (*16*). Our previous experience using NMP provided the rationale for a randomised controlled trial to assess its efficacy in preventing DGF in DCD kidneys (*17*), but the mechanisms by which NMP might benefit transplant kidneys are not fully understood. Furthermore, the question of whether additional manipulation of the kidney during NMP, for example by removal of pro-inflammatory cytokines and chemokines from the perfusate, might offer additional benefits in optimising the organ prior to transplantation has not been addressed in human kidneys, but our study in a porcine NMP model demonstrated promising results (*18*).

Here we took an unbiased approach to address these two questions using transcriptomic analysis of human kidney biopsies taken at the start and end of NMP to assess global changes in gene expression. We took advantage of the fact that we were able to access pairs of human kidneys from a single donor for research. The use of these kidney pairs allowed us to compare the effect of different interventions in two organs with an identical genetic background. We found that cold storage significantly reduced the expression of genes in key metabolic pathways, including oxidative phosphorylation (OXPHOS), compared with kidneys undergoing NMP. Following NMP, OXPHOS genes were upregulated, as were a number of immune and inflammatory pathway genes. This induction of immune genes was attenuated by the addition of a haemoadsorber (HA) to the perfusion circuit. Haemoadsorption not only reduced inflammatory gene expression, but also increased OXPHOS pathway genes, suggesting that it may be a useful intervention that may improve organ viability and reduce immunogenicity.

## Results

### NMP results in an increase in OXPHOS and inflammatory pathway genes compared with cold storage

To investigate the potential mechanisms by which NMP might impact kidney transplants, we took five kidney pairs, (n=4 DCD donors and n=1 DBD donor), that had a mean exposure to cold ischaemia of 1180.4 minutes (**Table 1**) and performed a time 0 (0hr) cortical biopsy prior to placing one of the pair back into cold storage and the other onto an NMP rig, as described previously (*10*) (**Figure 1A**). After 2 hours, a second biopsy was taken and RNA sequencing (RNA-Seq) performed on both samples. When comparing gene expression between the time 0 and 2 hour biopsies, we found that kidneys stored in cold storage had no statistically significant change in the expression of any individual gene when corrected for multiple testing (**Figure 1B**, left panel). In contrast, during the course of 2 hours of NMP, 956 genes were up-regulated and 353 genes were down-regulated (**Figure 1B**, right panel). We next assessed changes the expression of groups of genes within a common pathway rather than individual genes, gaining statistical power and identifying potentially important functional processes. Gene set enrichment analysis demonstrated that cold storage had a substantial impact on the expression of genes in a number of metabolic pathways (**Figure 1C**, left panel). In particular, there was a marked reduction in genes involved in OXPHOS, a key pathway required to generate ATP (*19*). In contrast, OXPHOS was among the pathways significantly up-regulated during NMP (**Figure 1C**, right panel), with potential benefits for cell viability and the restoration of cellular homeostasis. In addition, a number of pathways involved in immune or inflammatory processes were induced during NMP, with ‘TNFα signalling via NFkB’ demonstrating the largest increase. In keeping with this, *TNF* was among the top 20 most upregulated genes in the 2 hour NMP biopsies (**Figure 1D**), which also included *IL1B* and the neutrophil-recruiting chemokines *CXCL8* (IL8) and *CXCL2*. String analysis of the top 50 upregulated genes revealed the ordered upregulation of biochemically related genes which were clustered into four major nodes; IL8 and neutrophil-recruiting chemokines, Inflammasome-associated genes, NFkB signalling, and transcriptional regulation (**Figure 1E**). Together, our analysis demonstrates that during NMP there is an increase in the expression of genes in pathways that promote the generation of energy, with potentially beneficial effects for the organ, but a simultaneous induction in pro-inflammatory genes, the effect of which may be deleterious.

**Table 1.** Donor demographic in paired kidney NMP vs CS study

|  | Donor information |
| --- | --- |
| **Donor Type** | DBD = 1, DCD = 4 |
| **Age** | Mean= 66.8, SD = 10.42 |
| **Gender** | Male = 3, Female = 2 |
| **CIT (min)** | Mean = 1180.4, SD = 418.54 |
| **Samples collected** | Pre, 2hr |

**Figure 1.**
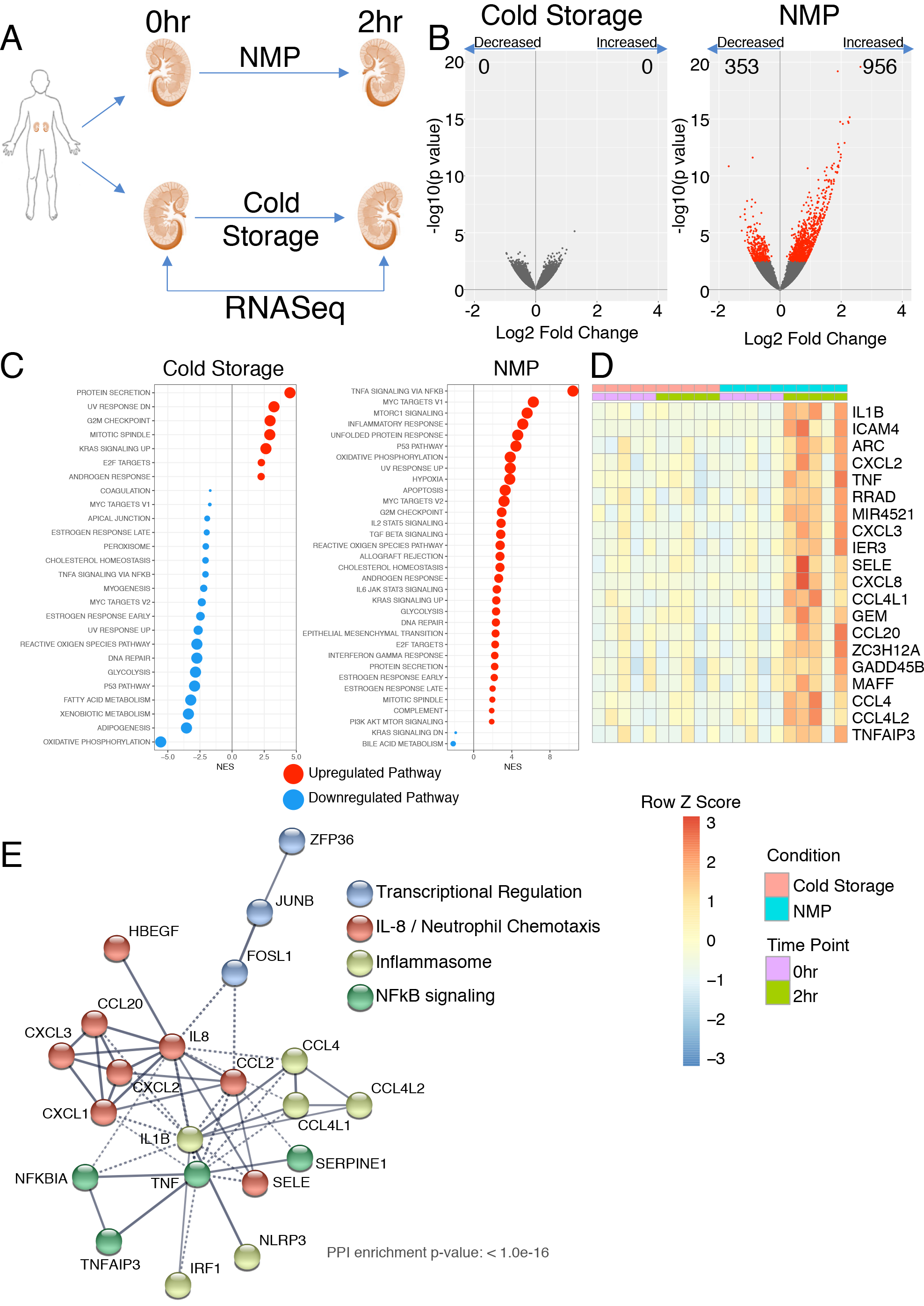
Kidneys exposed to cold storage show limited changes in gene expression compared with those undergoing NMP. **A –** Pairs of kidneys were obtained which had been declined for use in transplantation. One kidney was left on ice and the other underwent normothermic machine perfusion (NMP). Biopsies were taken from the outer cortex at the start and after 2hrs. **B** – Volcano plot indicating change in gene expression at 2 hrs for the indicated group compared to the start. Red dots indicate differentially expressed genes with an adjusted p value < 0.05 and the experimental group is indicated above the plot. **C** Gene set enrichment analyses of the differential expressions from B against the hallmarks pathways. Only significant pathways are plotted. Red dots indicate positive enrichment and blue negative, the size of the dot is inversely correlated with the FDR q value and the position indicates the normalised enrichment score (NES). **D** – Heatmap of the top 20 significantly unregulated genes during NMP, genes are ranked by log 2 fold change. **E** – STRING analysis of the top 50 genes upregulated during NMP. The colour of each node indicates membership of each cluster.

### Urine output and renal blood flow during NMP demonstrate differing associations with OXPHOS and inflammatory pathway genes

The quantity of urine produced during NMP is one of a number of parameters included in surgical quality assessment scores used to guide organ utilisation decisions (*15*), but whether high urine output during NMP truly portends a good prognosis for the kidney and the underlying molecular pathways activated in kidneys with a high urine output is unclear. In n=10 kidney undergoing NMP (NMP only kidneys, **Table 1** **and** **2**), including the 5 described in Figure 1 above, we observed a range of urine outputs from 0 to 340ml over the 2 hour period of perfusion (**Figure 2A**). Kidneys with a higher urine output demonstrated differential expression of 11 genes, including heat shock proteins (HSPs), HSPA1A, HSPA1B and HSPH1 (**Figure 2B**). HSPs are protein chaperones and form part of the canonical cellular response to a variety of physiological stressors (*20*) and have been shown to play both beneficial and detrimental roles in murine models of kidney disease (*21*). In models of ischaemic renal injury, HSP deficiency attenuated disease severity, an effect thought to be mediated by the pro-inflammatory effects of HSPs which act as potent stimulators of innate immunity when released extracellularly (*22*). Gene set enrichment analysis showed that urine output positively correlated with the ‘TNFα signalling via NFkB’ pathway genes, and negatively correlated with OXPHOS genes (**Figure 2C-D**, **S1**). Similarly, in pre perfusion biopsies, OXPHOS negatively correlated with urine output in these kidneys, whilst pathways associated with immune activation including ‘TNFα signalling via NFkB’ and ‘allograft rejection’ were positively correlated with urine output, suggesting that NMP had little effect on these processes in high urine output kidneys (**Figure 2C-D**, **S1**). These data challenge the dogma that urine output occurs in more viable, ‘healthy’ kidneys and suggest that in fact these kidneys may have more inflammatory potential, are less able to generate energy and that this is not substantially altered during NMP.

**Table 2.** Donor demographic in paired kidney NMP vs NMP + HA study

|  | Donor information |
| --- | --- |
| **Donor Type** | DBD = 1, DCD = 4 |
| **Age** | Mean= 65.2, SD = 3.70 |
| **Gender** | Male = 5, Female = 0 |
| **CIT (min)** | Mean = 1447.2, SD = 500.98 |
| **Samples collected** | Pre, 2hr, 4hr |

**Figure 2.**
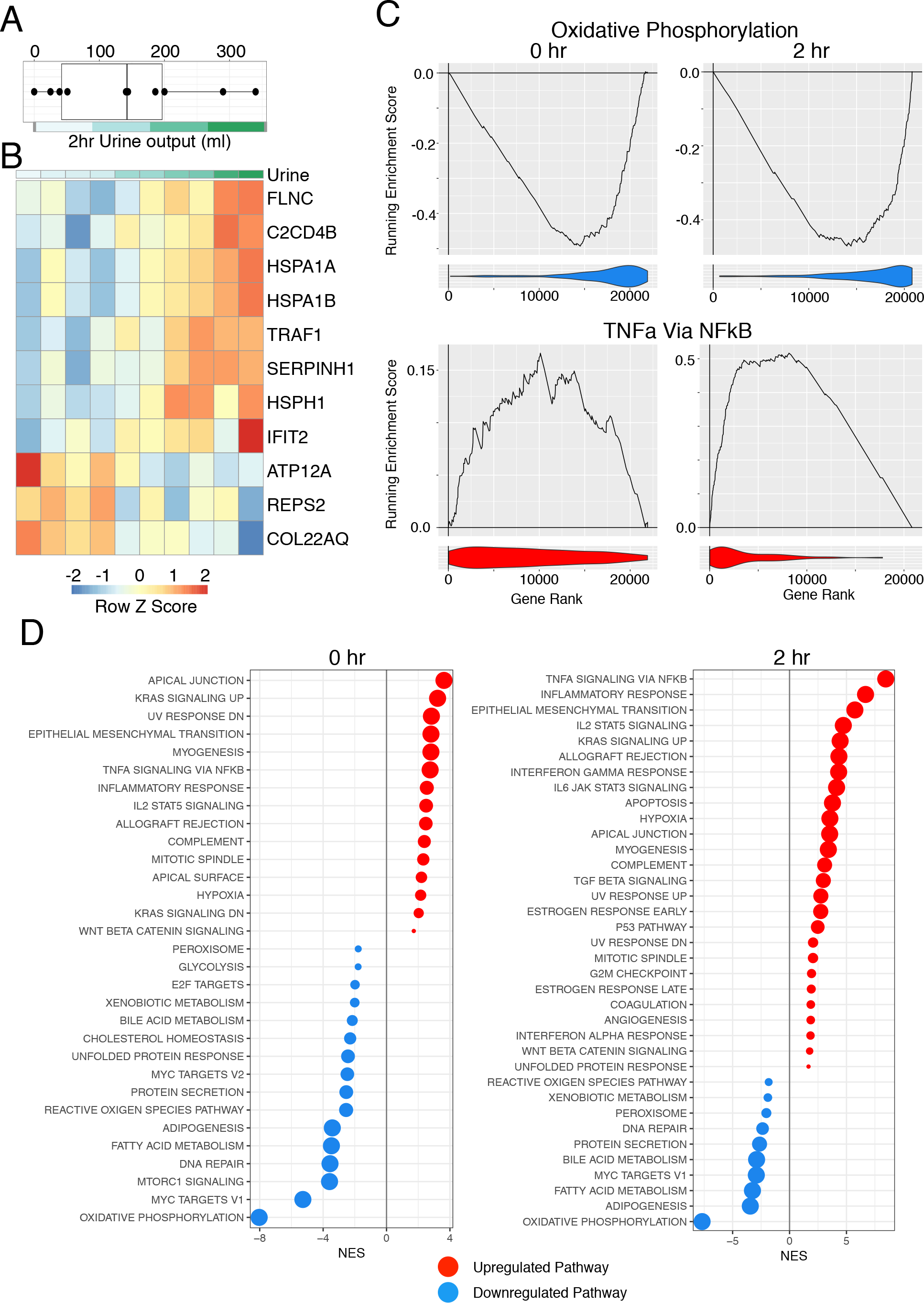
Correlation of transcriptome of NMP after 2 hrs with urine output. **A** – Total urine output by each kidney after 2 hrs Normothermic machine perfusion (NMP). **B** – Heatmap of all genes that are significantly correlated with urine output in biopsies taken post NMP. **C** – Enrichment plots from GSEA for key pathways from the Hallmark database. Analysis is for the correlation of urine output after 2 hours with the transcriptome of the kidney either pre (0hr) or post 2hr NMP. The line indicates the running enrichment score and the violin plot indicates the distribution of the member genes of the geneset throughout the ranked gene list used in each analysis. **D** GSEA for the analysis from A against the hallmarks database of genesets. Only significant pathways are plotted. Red dots indicate positive enrichment and blue negative, the size of the dot is inversely correlated with the FDR q value and the position indicates the normalised enrichment score (NES).

Renal blood flow during perfusion has also been assessed as a parameter that may reflect subsequent transplant function (*15*). In the 10 kidneys studied, renal blood flow varied from 14.1-168.8 ml/minute (**Figure 3A**). 8 genes were significantly positively correlated in kidneys with renal blood flow (**Figure 3B**). Gene set enrichment analysis demonstrated that high renal blood flow positively correlated with OXPHOS genes in 0 hour biopsies, but in contrast to those with the highest urine output, perfusion had a significant impact, resulting in a negative correlation with ‘OXPHOS’ pathway genes by 2 hours (**Figure 3C-D**, **S2A**). Renal blood flow negatively correlated with a number of immune and inflammatory gene pathways in both 0 and 2 hour biopsies, in contrast to urine output (**Figure 3C-D**, **S2B**). Together these data suggest that renal blood flow and urine output may not be equivalent, generic indicators of more viable kidneys but indicate different underlying processes may be occurring in these kidneys.

**Figure 3.**
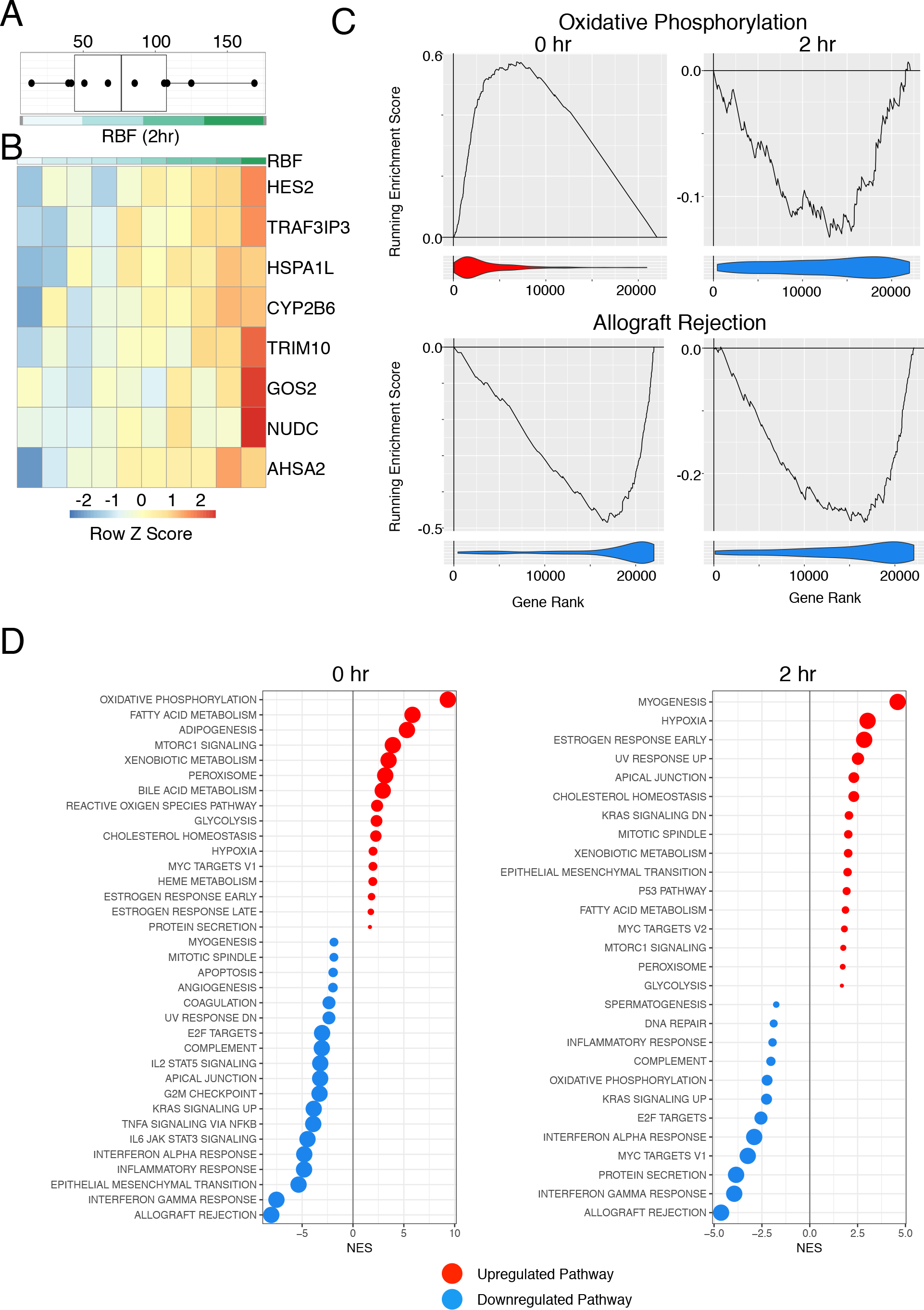
Correlation of the transcriptome with renal blood flow. **A** – Renal blood flow after 2 hours of normothermic machine perfusion (NMP). **B** – Heatmap of all genes which correlate with 2hr RBF in samples taken after 2hrs NMP.**C** - Enrichment plots from GSEA for key pathways from the Hallmark database. Analysis is for the correlation of RBF at 2 hours with the transcriptome of the kidney either pre (0hr) or post 2hr NMP. The line indicates the running enrichment score and the violin plot indicates the distribution of the member genes of the geneset throughout the ranked gene list used in each analysis. **D** GSEA for the analysis from A against the hallmarks database of genesets. Only significant pathways are plotted. Red dots indicate positive enrichment and blue negative, the size of the dot is inversely correlated with the FDR q value and the position indicates the normalised enrichment score (NES)

### Addition of a haemoadsorber to the perfusion circuit has no effect on urine output and renal blood flow

Analyses of kidney perfusates have demonstrated a substantial increase in the concentration of a number of pro-inflammatory cytokines and chemokines during the course of hypothermic and normothermic machine perfusion (*23, 24*). These bioactive molecules re-circulate into the kidney, with the potential to further induce inflammation in the vasculature and beyond. We therefore reasoned that the addition of a haemoadsorber (HA) to the NMP circuit to remove cytokines and chemokines may have beneficial effects for the organ. Such approaches have shown some efficacy in the treatment of patients with systemic inflammatory response syndrome (*25, 26*), and in porcine kidneys undergoing NMP, this strategy increased renal blood flow (*18*). To test the effect of cytokine and chemokine adsorption in human kidneys, we took 5 kidney pairs and performed NMP for 4 hours. In each case, a Cytosorb haemoadsorber that removes molecules with a molecular weight of 10-50kDa was added to the perfusion circuit of one of the kidney pairs (NMP+HA) (**Figure 4A**). We confirmed that in the presence of this HA, there was a reduction in a number of cytokines, including IL1β and IL6 (**Figure 4B**). However, the addition of the HA had no effect on renal blood flow or urine output compared with kidneys undergoing NMP alone (**Figure 4C, D**). Oxygen consumption and acid-base homeostasis were also similar between kidney pairs and were not significantly impacted by the addition of the HA (**Table 3**). Thus, over 4 hours of NMP, the HA had no impact on the parameters currently used clinically to generate quality assessment scores.

**Table 3.** Levels of pH, bicarbonate, base excess and oxygen consumption during normothermic machine perfusion (NMP)

|  | NMP |  |  | NMP+HA |  |  |
| --- | --- | --- | --- | --- | --- | --- |
|  | Pre | 1h | 4h | Pre | 1h | 4h |
| **pH** | 7.29 ± 0.1 | 7.40 ± 0.1 | 7.47 ± 0.1 | 7.26 ± 0.1 | 7.36 ± 0.1 | 7.43 ± 0.1 |
| **Bicarbonate (mmol/L)** | 38.7 ± 12.5 | 28.5 ± 5.7 | 21.0 ± 3.6 | 31.6 ± 7.4 | 26.0 ± 6.2 | 24.2 ± 8.2 |
| **Base excess (mmol/L)** | 9.8 ± 12.3 | 3.3 ± 6.8 | -1.9 ± 4.0 | 3.0 ± 7.1 | 0.2 ± 6.2 | -0.55 ± 10.0 |
| **Oxygen consumption (ml/min/100g)** |  | 1.19 ± 0.10 | 1.24 ± 0.16 |  | 1.41 ± 0.24 | 1.27 ± 0.16 |

**Figure 4.**
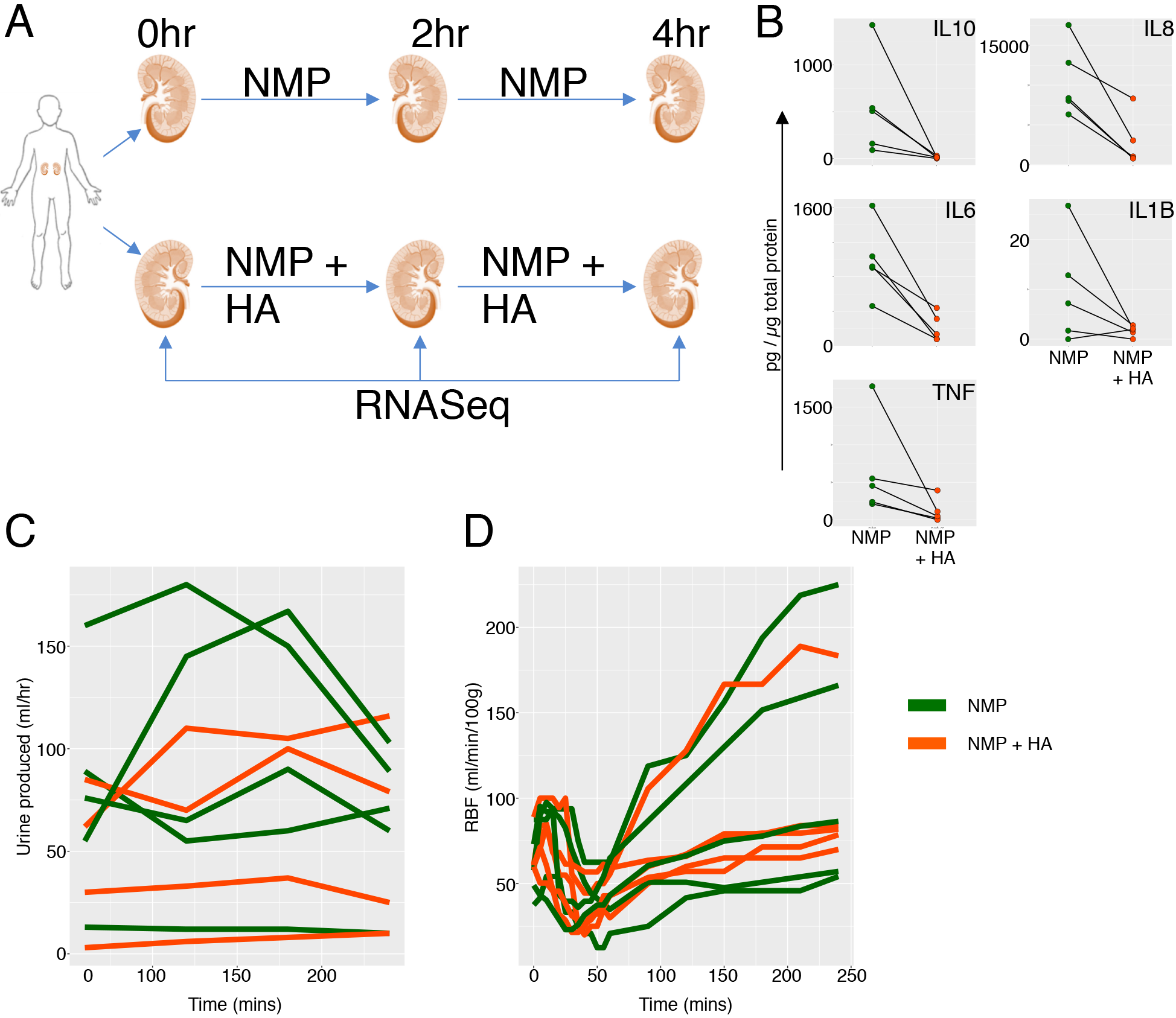
Addition of a haemoadsorber to the Normothermic machine perfusion circuit has substantial effects on cytokine level in perfusate but not on physical parameters of rig. **A –** Pairs of human kidneys were taken from the same donor, one of the pair underwent the standard normothermic machine perfusion (NMP) protocol for 4 hours and the other kidney underwent NMP with the addition of a haemoadsorber to the circuit. Samples were taken for RNA-Seq prior to perfusion (0hr), after 2 hrs and at the end (4hrs). **B** – Concentration of key cytokines in the perfusate after 4hrs NMP. Line indicates pairs of kidneys. Cytokine measurements were normalised to total perfusate protein content and are measured in pg/*μ*g of total protein. **C** & **D** – Urine production and renal blood flow (RBF) across the timecourse of perfusion. Green line indicates NMP alone and orange with the addition of the heamoadsorber.

### Addition of a haemoadsorber to the perfusion circuit has a substantial impact on gene expression, reducing inflammatory pathway genes and increasing OXPHOS pathway genes

Despite the lack of effect on perfusion parameters, we found a substantial effect of the HA on gene expression. At 2 hours post-perfusion, n=1794 genes were upregulated in NMP kidneys but only half this number (n=898) were increased when the HA was present (**Figure 5A**). Similarly, at 4 hours n=4026 genes were up-regulated in the NMP group and n=2606 in the NMP+HA group (**Figure 5A**). The number of genes down-regulated was also reduced by the addition of HA (**Figure 5A**). Of note, n=1702 genes were upregulated at both 2 and 4 hour timepoints (**Figure 5A**), including TNF and *IL6* (**Figure 5B**). The addition of a HA reduced the up-regulation of these pro-inflammatory genes (**Figure 5B**), supporting the thesis that re-circulating small molecules can exacerbate inflammation in perfused organs. Further differences were observed between the groups for many members of the cytokine and chemokine gene families (**Figure S3A and B**). To address difference in gene expression as a result of the addition of a HA to NMP, we directly compared the experimental groups after 4 hours of NMP and found 46 genes were significantly upregulated and 181 downregulated with the addition of the HA (**Figure 5C**). In keeping with this, gene set enrichment analysis showed a significant decrease in the ‘TNFα signalling via NFkB’ pathway in NMP+HA kidneys compared with NMP alone (**Figure 5D, E**). However when the level of TNF protein was measured in the tissue post EVLP no difference was observed suggesting that differences in the transcriptome had not yet been translated to the proteome at this early time-point (**Figure S3C**). Notably, the presence of the HA not only reduced inflammatory gene expression within the kidney but also increased OXPHOS pathway and fatty acid metabolism pathway genes, both of which contribute to energy generation (**Figure 5D, E**). This demonstrates that soluble mediators released from the kidney recirculate and drive *de novo* expression of inflammatory genes within the organ, and potentially have additional deleterious effects on energy generation.

**Figure 5.**
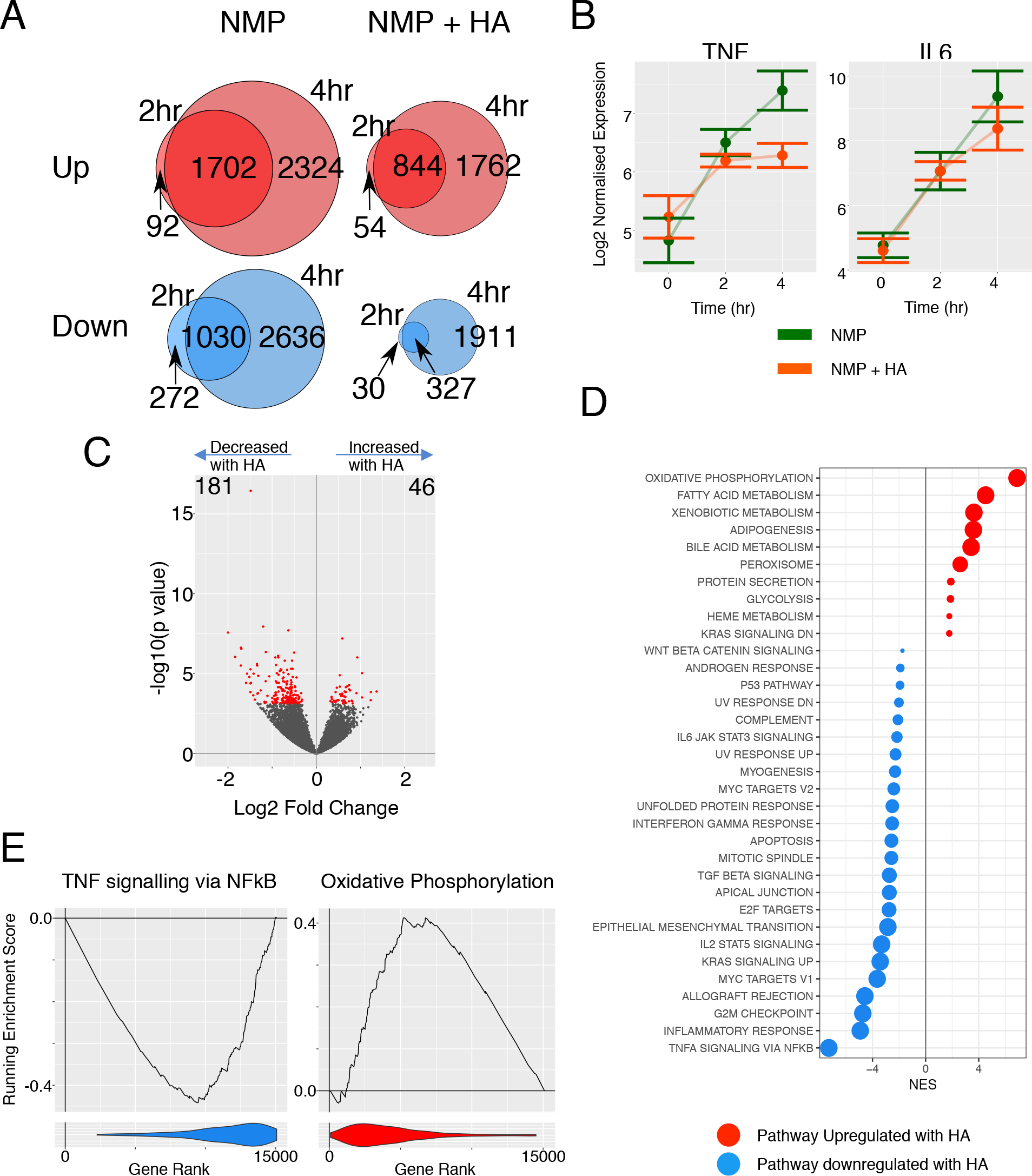
Addition of a haemoadsorber to the NMP circuit has substantial effects on gene expression. **A** – Venn diagram showing the numbers of genes significantly differently expressed when comparing the 2 hr or the 4hr samples to the pre (0hr) samples for the indicated groups. Red diagrams are upregulated genes and blue downregulated. The intersection of the sets in each venn diagram are the genes which are deferentially expressed at both time points in the same direction. **B** – log2 normalised expression values for the indicated genes across perfusion. Green line indicates NMP alone and orange with the addition of the heamoadsorber (HA). **C** – Volcano plot for pairwise comparison of NMP alone with NMP + HA at 4hrs relative to NMP alone. Red dots indicate differentially expressed genes with an adjusted p value < 0.05. **D** – GSEA analysis of the results from C against the hallmarks pathway of genesets. Only significant pathways are plotted. Red dots indicate positive enrichment and blue negative, the size of the dot is inversely correlated with the FDR q value and the position indicates the normalised enrichment score (NES). **E** Enrichment plots from GSEA for key pathways from the Hallmark database from D. The line indicates the running enrichment score and the violin plot indicates the distribution of the member genes of the geneset throughout the ranked gene list used in each analysis.

### Haemoadsorption decreases NLRP3 inflammasome activation and the generation of IL1β

To further assess the impact of HA on the induction of sterile inflammation in the kidney, we analysed genes associated with NLRP3 inflammasome activation. This showed a reduction in the expression of *IL1B*, *NLRP3*, and *CASP1* (**Figure 6A, B).** Animal models suggest that neutrophil recruitment to injured kidney may cause collateral damage and increase kidney injury (*27, 28*). The impact of the HA on the expression of neutrophil-recruiting chemokines varied, with all five kidneys showing a reduction in *CXCL8* and *CXCL2* but the magnitude of this decrease was substantial in three kidneys (**Figure 6C, D**).

**Figure 6.**
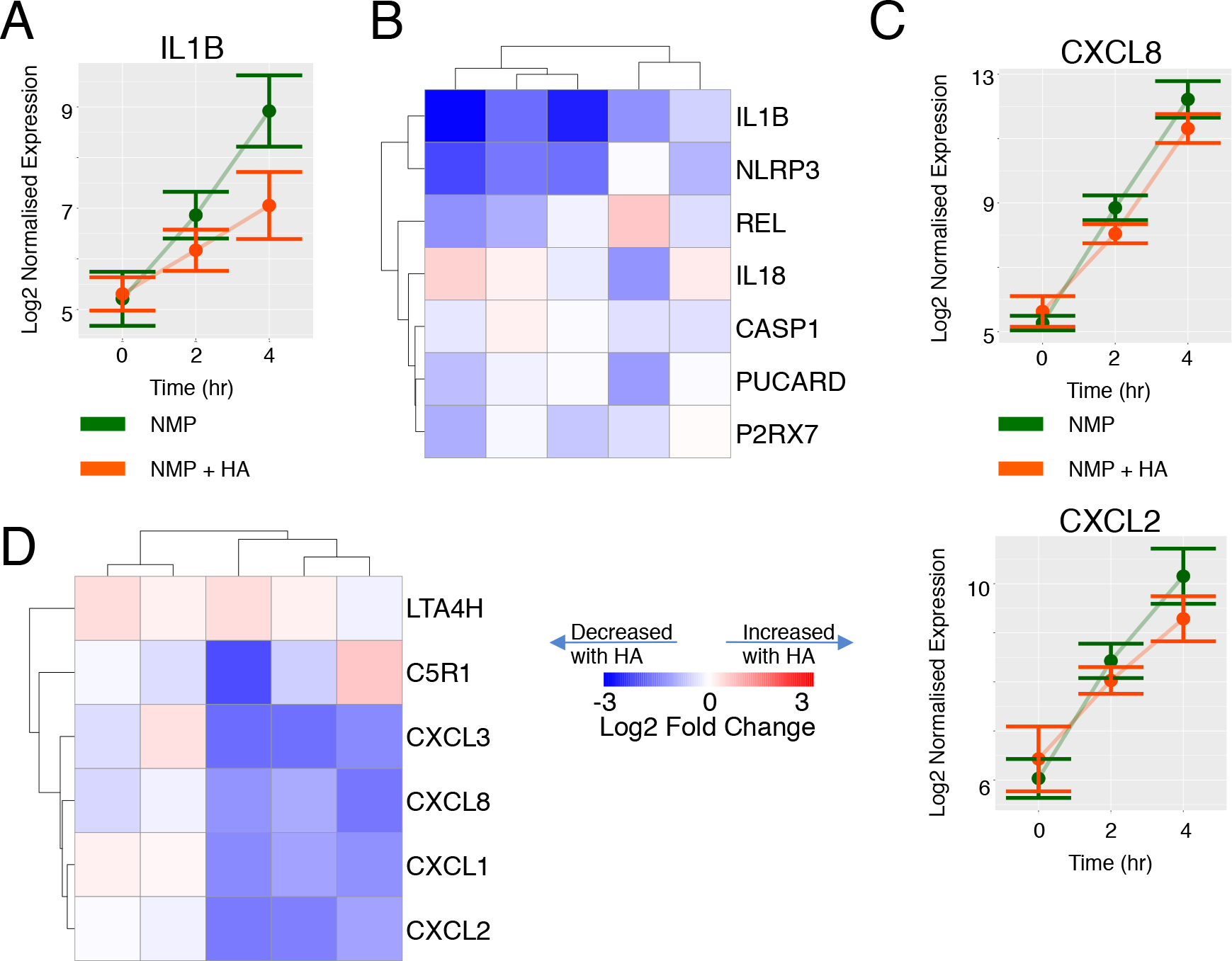
Effect of haemoadsorber filter on inflammasome activation and neutrophil modulation during NMP. **A** – Log2 normalised expression values for IL1B across perfusion. Green line indicates normothermic machine perfusion (NMP) alone and orange with the addition of the heamoadsorber (HA). **B** Heatmap of log2 fold change of inflammasome related genes comparing kidneys which have undergone NMP with a HA filter to NMP alone. Each column represents the comparision of 1 kidney to its pair. **C** Expression of the indicated genes as for A. **D** Heatmap showing Log2 fold change in neutrophil recruiting chemokines comparing kidneys which have undergone NMP with a HA filter to NMP alone.

Overall, the transcriptional changes we have identified suggest that NMP has potential benefits over cold storage in terms of its effects on energy generation in the kidney. However, during perfusion, some bioactive molecules are released from the kidney into the perfusion circuit, generating an amplification loop that drives inflammation when they re-enter the kidney. Removal of these molecules interrupts this loop, and may be useful to reduce inflammation and increase energy generation, further enhancing the beneficial effects of NMP (**Figure 7**).

**Figure 7.**
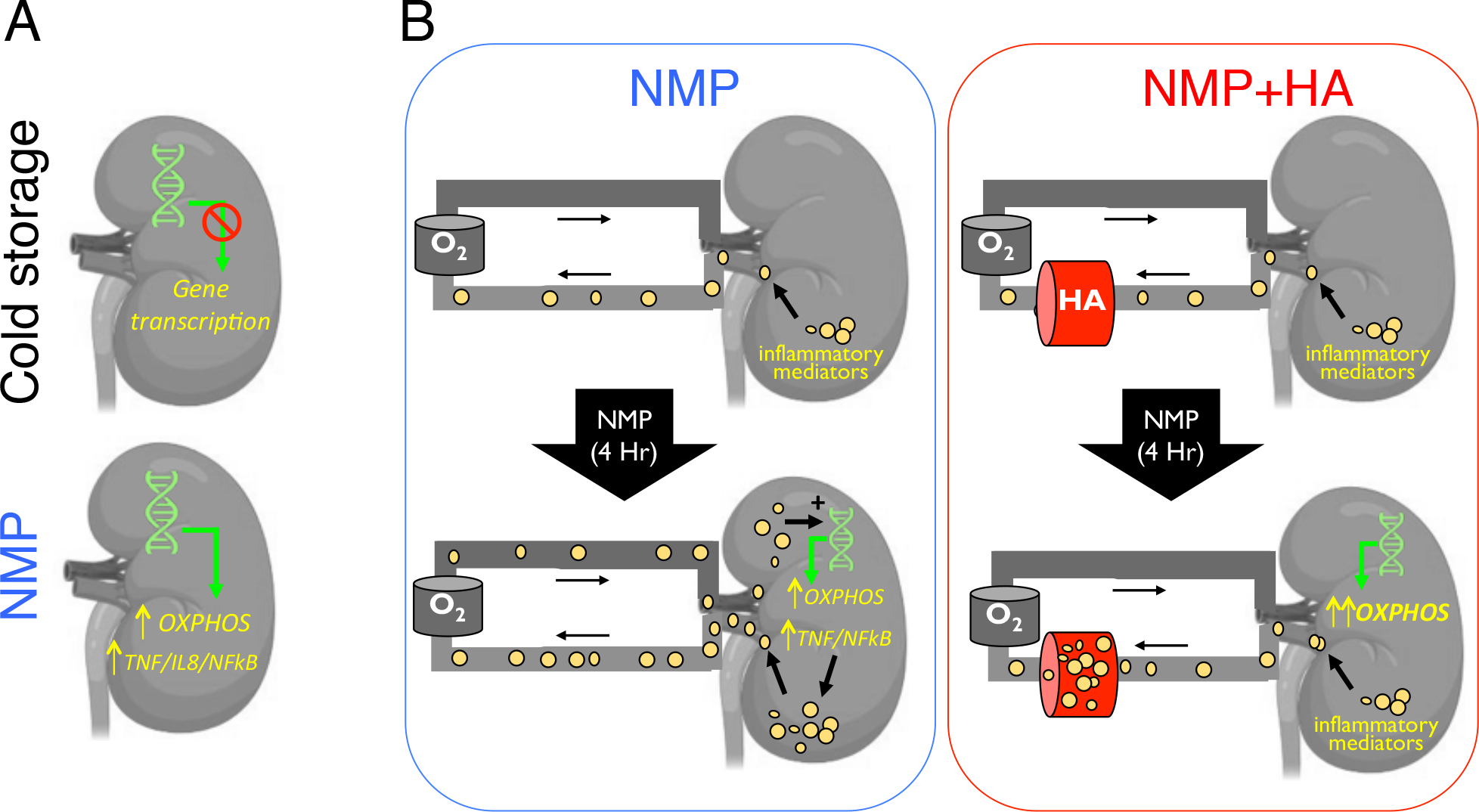
Diagrammatic summary. **A** During cold storage there is a global reduction in transcription. During NMP the expression of genes in a number of pathways are upregulated including oxidative phosphorylation (OXPHOS) and inflammatory pathway genes such as *TNF*, *IL8* and *NFkB*. **B. Left panel** – During NMP inflammatory mediators (yellow circles) are released from the kidney into the perfusion solution. They re-circulate and stimulate pro-inflammatory gene transcription in the kidney, and are associated with a reduction in energy pathway production genes, reducing ATP. **Right panel** - The presence of the haemoadorber (HA) breaks this inflammatory amplification loop

## Discussion

Our data show that cold storage is effective in limiting substantial changes in gene expression. This is in-keeping with rodent models, where kidneys stored for up to 18 hours in cold storage showed little change in the expression of pro-inflammatory cytokines, including IL1β, TNF and IL6 (*29*). Our use of unbiased, global transcriptional profiling rather than the measurement of a small number of candidate genes allowed us to analyse the expression of groups of genes found within specific pathways. This revealed a significant reduction in the expression of OXPHOS and glycolysis pathway genes in cold stored kidneys, potentially reducing the capacity of these kidneys to generate ATP. This observation is in keeping with a previous study showing reduced ATP levels in human kidneys following cold storage (*30*). A number of pro-inflammatory pathways were also down regulated during cold storage, including TNFα activation via NFkB and reactive oxygen species pathways which may be potentially beneficial in reducing sterile inflammation in the kidney following transplantation. However, it may be that cold storage merely puts a temporary hold on these pathways, and that following re-perfusion in the recipient similar changes in inflammatory gene expression as observed in NMP would occur.

NMP had the opposite effect to cold storage on OXPHOS and glycolysis pathway genes, increasing their expression, with the potential to increase the cellular capacity to generate ATP and to restore homeostasis. Of note, hypothermic oxygenated perfusion of human kidneys has also been shown to increase ATP levels compared with cold storage (*30*), therefore the restoration of oxygenation may be the principle stimulus for these processes rather than normothermia. Nonetheless, these changes are likely to be beneficial for the organ.

We also assessed the molecular processes that correlate with urine output and renal blood during NMP. These parameters have previously been used, along with a number of other measures, to generate a quality assessment score of perfused kidneys. High values of urine output and renal blood flow have been considered to reflect a more viable graft (*15*), but whether they provide a similar generic readout of a more healthy kidney, or potential reflect different underlying molecular pathways that are activated in kidneys is unclear. Our data reveal that pathways correlating with high urine output and high renal blood flow differ, and in fact, these parameters demonstrate polar opposite associations with inflammatory pathways. High urine output was associated with higher expression of immune pathway genes whilst high renal blood flow was negatively correlated with these pathways. Metabolically, higher urine output correlated with lower expression of OXPHOS pathway genes both prior to NMP and after 2 hours. In contrast, higher renal blood occurred in kidneys with high OXPHOS pathway gene prior to NMP. Overall, our data would suggest that renal blood flow may be a more useful indicator of graft functionality, but the true prognostic significance of these parameters will need to be confirmed in large prospective clinical studies, such as the one we are currently undertaking (*17*).

During 4 hours of NMP there was an induction of inflammatory genes in the kidney. However, it is worth noting that current clinical practice involves 1 hour of NMP only, and this effect may not be evident over a shorter perfusion time. However, longer perfusion may have benefits in terms of restoration of oxygenation and energy generation, and in our experiment, the negative effects of immune gene induction could be substantially negated by the introduction of a cytosorb HA to the perfusion circuit. This device removes molecules with a molecular weight of 10-50kDa, including cytokines and chemokines and many other bioactive molecules. Indeed, we observed a decrease in several cytokines including IL1β, TNF◻ and IL6, although the reduction was not statistically significant due to the modest sample size. Perfusion parameters were not affected by the addition of the HA, but haemoadorption had a substantial impact on the transcriptome of the kidney after 4 hours of NMP. Overall, it halved the number of genes that were differentially expressed during NMP. This included a marked attenuation in the induction of pro-inflammatory immune pathway genes and in NLRP3 inflammasome-associated genes. Our data suggest that inflammatory mediators generated by the kidney during the course of NMP enter the perfusion circuit and are capable of exacerbating sterile inflammation, and indicate that their removal would be beneficial to negate the marked induction of inflammation pathway genes observed during NMP. The addition of the HA to the NMP circuit also had potential beneficial effects on metabolic pathways, increasing the expression of both OXPHOS pathway and fatty acid metabolism pathway genes. Fatty acid catabolism provides an important source of energy, providing a substrate (acetyl CoA) for the citric acid cycle to generate ATP (*31*). Together, the induction of these pathways has the potential to reduce the tubular damage occurring during the process of transplantation. Indeed, a reduction in fatty acid oxidation (FAO) gene expression in renal tubular cells has been observed in acute kidney injury and restoration of renal FAO gene expression with peroxisome proliferator activated receptor-alpha ligands ameliorated injury (*32*).

There are a number of limitations to our study. Firstly, although the kidney pairs used were from the same individual and therefore genetically identical, there may still be asymmetrically affected by pathology, for example, cysts, pyelonephritis or renal arterial disease leading to variable susceptibility to the effects of the interventions investigated. Secondly, although we assessed the impact of HA on gene expression, the kidneys we studied were not subsequently transplanted. Therefore were unable to directly correlate changes in gene expression observed with cold storage, NMP or NMP+HA with subsequent clinical outcomes.

In summary, our study provides the first global transcriptional profile of human kidneys undergoing NMP, resolving the differing molecular pathways that are activated in NMP compared with cold storage, and showing that the deleterious effects of bioactive molecules produced or released from the kidney during NMP can be reversed by the addition of a haemoadsober. The results provide the mechanistic foundation for applying such an intervention in the context of a clinical trial. This will be required to determine if the attenuation in pro-inflammatory gene expression and increase in ATP generating metabolic pathway genes we have observed has a significant impact on clinical parameters such as delayed graft function. Our data also has implications for perfusion strategies beyond the kidney, including in liver and lung transplantation, suggesting that the removal of bioactive molecules from perfusates should be investigated in these contexts where NMP is increasingly used. Finally, our study highlights the utility of global transcriptional profiling for assessing novel interventions to perfused organs; transcriptional changes precede changes in protein abundance (traditionally used as biomarkers of kidney injury) and tens of thousands of gene transcripts can be readily measured. Thus, mRNA measurement has the potential to provide an early, sensitive readout of cellular function of human organs retrieved for transplantation that can be applied to future studies.

## Methods

### Study design

Ethical approval was granted from the national ethics committee in the UK REC reference (15/NE/0408). 10 pairs of human kidneys rejected for transplantation were recruited into the study. The study was divided into two parts. In the first study five pairs of kidney were randomly assigned to either cold storage or normothermic machine perfusion (NMP) for 2h.

In the second study five pairs of kidneys were randomly assigned to either NMP with (NMP+HA) or without (NMP) the addition of a haemoadsorber (Cytosorb, Linc Medical, Leicester, UK) in the perfusion circuit (18). Kidneys were perfused for 4h.

### Normothermic machine perfusion

After a period of cold storage the kidneys were weighed and prepared for perfusion. The renal artery, vein and ureter were cannulated and kidneys flushed with 1L of cold Ringer’s solution to remove the preservation solution. Perfusion was carried out using an adapted paediatric cardiac bypass system (Medtronic, Bioconsole 560) as described previously (10). The perfusion system was primed with 300ml Ringer’s solution (Baxter Healthcare, Thetford UK), 15ml Mannitol 10% (Baxter Healthcare), 27 ml sodium bicarbonate 8.4% (Fresenius Kabi, Runcorn, UK), 3000iu heparin (LEO Pharma A/S, Ballerup, Denmark) and 6.6mg Dexamethasone (Hameln Pharmaceuticals, Hamelin, Germany). A unit of compatible packed red cells was then added. The red cell-based solution was oxygenated with a balance of 95% oxygen/5% CO_2_ at a flow rate of 0.1L/min and warmed to 35.5 - 36.5°C.

The red cell-based solution was circulated continually through the kidney via the renal artery at a mean arterial pressure of 85mmHg and pump speed of 1450RPM. A nutrient solution (Synthamin 17 10%, Baxter Healthcare, Thetford, UK) with 15ml of sodium bicarbonate 8.4% (B Braun, Melsungen, Germany) and 100IU of insulin added (Actrapid, Novo Nordisk, London, UK) was infused at a rate of 20ml/h, glucose 5% (Baxter Healthcare) at a rate of 5ml/h and Ringer’s solution was used to replace urine output (ml for ml). Epoprostenol sodium 0.5mg (Folan, Glaxo Wellcome UK Ltd, Uxbridge, UK) was infused at rate of 5ml/h throughout perfusion to enhance blood flow.

### Normothermic Machine Perfusion Outcome Measures

The renal blood flow (RBF) was recorded every 5min for the 30min and thereafter every 30min. Samples of perfusate were collected pre-perfusion and after each hour of perfusion for analysis of haematology and urea & electrolytes (U&Es). Urine samples were also collected hourly for the measurement of U&Es. Samples of perfusate were also retained for the measurement of cytokines. Perfusate was centrifuged at 1600rpm for 10min at 4°C. The supernatant was collected and frozen in liquid nitrogen then stored at −70°C until analysed.

Samples of arterial and venous perfusate were collected at 1 and 4h of perfusion for blood gas analysis (OPTI-CCS, Una Health, Stoke-on-Trent, UK). Oxygen consumption was calculated using the following equation;

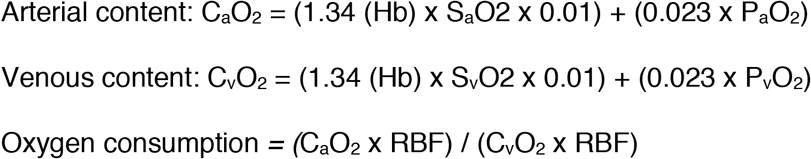

Core biopsies were taken from each kidney at time 0 and after 2h of cold storage or perfusion in the first study and at time 0, 2h and 4h of perfusion in haemoadsorber study. Biopsies were stored in RNAlater solution (Invitrogen RNAlater™Soln.) for fixing.

### RNA extraction

RNA was extracted from biopsies stored in RNALater (Ambion) at −80C. Biopsies were removed from their storage solution and placed with 1ml Lysis Buffer (Ambion) in a MK28-R grinder tube (Bertin Instruments) and lysed using a Precellys 24 homogeniser (Bertin Instruments). Tubes were subsequently centrifuged at 1500xg for 4 minutes, the supernatant removed and the RNA extraction performed using a pure link RNA mini kit (Ambion) as per manufacturers instructions. Contaminating DNA was removed using TURBO DNase (Ambion) as per manufacturers instructions. Concentration of RNA was assessed using a Nanodrop Spectrophotometer (Thermo Scientific). Quality of RNA was assessed using a RNA nano Bioanalyzer kit (Agilent) using a Bioanalyzer 2100 (Agilent).

### RNA sequencing

0.5*μ*g of RNA was used for producing libraries for sequencing using TruSeq Stranded total RNA library prep kit (Illumina) as per manufactures instructions with a final PCR amplification of 14 cycles. Libraries were then sequenced on a Hiseq 4000 sequencer (Ilumina) by Genewiz.

### RNA sequencing analysis

Following sequencing data was demultiplexed to give individual fastq files using Casava (Illumina). Fastq files were assessed for quality control purpose using FASTQC. The Fastq files were aligned to the human genome (Hg38) using Hisat2(*33*). All further analysis was carried out using the R statistical environment. A table of gene counts was produced using the featureCounts function within Rsubread and normalisation and differential gene expression analysis was carried out using DESeq2. For GSEA genes were ranked by the inverse of the p value with the sign of the log fold change and then ran against the hallmarks database within MSigDB using the GSEA program from the broad with the pre ranked option.

### String analysis

The top 50 significant genes, ranked by log fold change for the effect of perfusion, were used to run STRING analysis. Gene names were converted into protein names and two proteins were considered connected if they had a mean interaction score of >0.7. Unconnected proteins were removed from the network and a force directed graph plotted. Network was subsequently clustered using k-means clustering and annotated using interpretation of GO terms associated with each cluster.

### Tissue protein extraction

Total protein was extracted from tissue by homogenisation, using a precellys (Bertin instruments), of a small biopsy (approx. 27mm^3^) in T-PER tissue protein extraction reagent (thermo fisher scientific) with HALT protease inhibitor (thermo fisher scientific). Protein content was measured using by BCA analysis (thermo fisher scientific) and samples normalised.

### Cytokine measurements

For all cytokine measurements analysis was carried out using DuoSet ELISA reagents (R&D) as per manufactures recommendations.

## Acknowledgments

**Funding:** J.R.F., S.H., and M.L.N. are supported by the NIHR Cambridge Blood and Transplant Research Unit Organ Donation. S.H., and M.L.N are also supported by the Stoneygate Trust (RG72312) T.M. was funded by an Evelyn Trust Research Grant (16/23). M.R.C. is supported by the National Institute of Health Research (NIHR) Cambridge Biomedical Research Centre, by a Medical Research Council New Investigator Research Grant (MR/N024907/1) and an NIHR Research Professorship (RP-2017-08-ST2-002). T.C.D. and C.J.W. by M.R.Cs Medical Research Council New Investigator Research Grant (MR/N024907/1). The research was funded by the National Institute for Health Research Blood and Transplant Research Unit (NIHR BTRU) in Organ Donation and Transplantation at the University of Cambridge in collaboration with Newcastle University and in partnership with NHS Blood and Transplant (NHSBT). The views expressed are those of the author(s) and not necessarily those of the NHS, the NIHR, the Department of Health or NHSBT.

## Author contributions

J.R.F., S.H., M.L.N, and M.R.C. designed the study and interpreted the data. J.R.F., S.H., T.M., C.J.W, and T.C.D. performed experiments, J.R.F., S.H., and M.R.C created figures and tables, M.R.C. wrote the main manuscript, J.R.F. and S.H. the methods and Figure legends, J.R.F, S.H. and M.L.N. edited the manuscript.

## Non-author contributions

The authors thank all organ donors and their families. The Clatworthy Lab are grateful for the core facilities provided by the MRC Laboratory of Molecular Biology.

