## Supplemental Figures 1-3 for "Investigation of the transcriptional profile of human kidneys during machine perfusion reveals potential benefits of haemoadsorption"

### Supplemental Fig 1

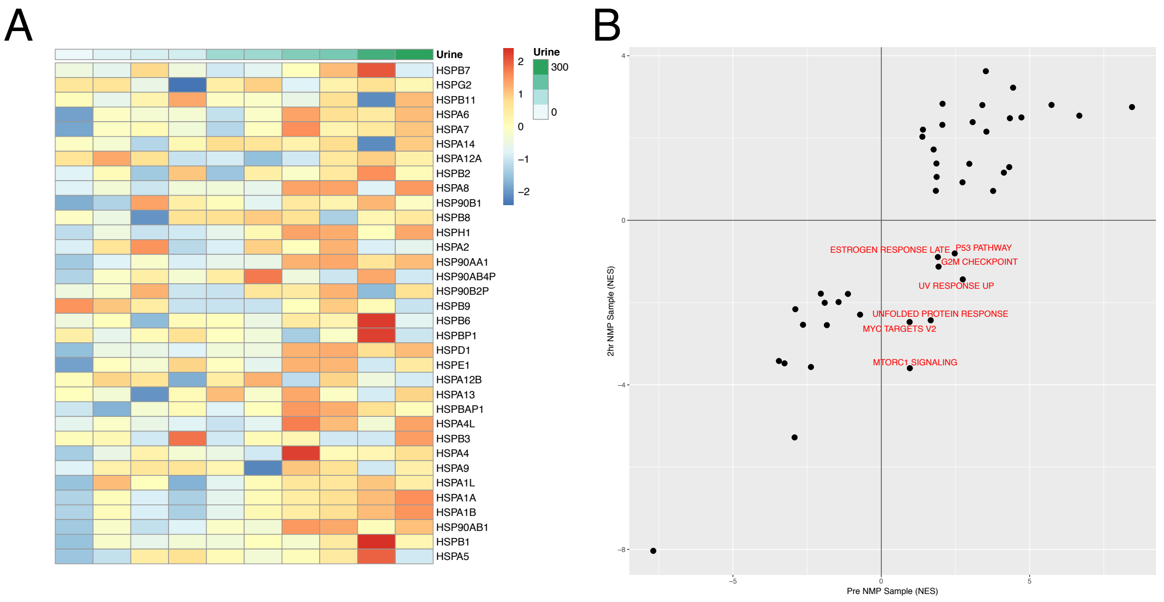


**Supplemental figure 1 - Correlation of transcriptome of kidneys prior to NMP with urine output after 2 hours. A** Heatmap of all HSP genes ordered by urine output. **B** Comparison of enrichment scores for correlation pre and post 2 hrs NMP with urine output by GSEA, against the Hallmarks database. Only pathways which are significant in at least 1 comparison have been plotted. Pathways which are altered between the conditions are labelled.

### Supplemental Figure 2

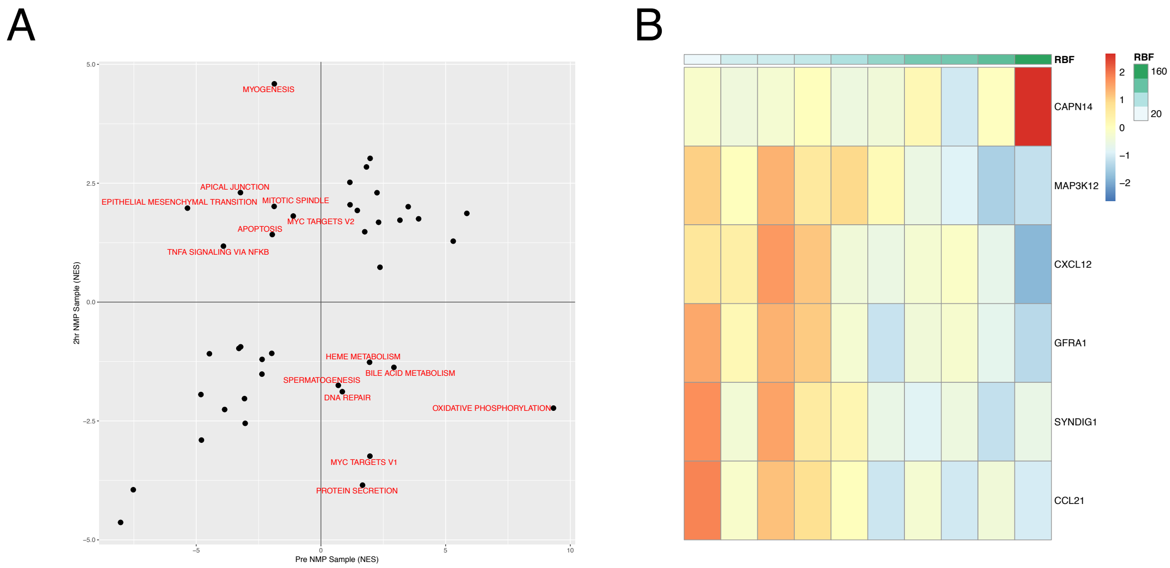

**Supplemental Figure 2 – Correlation of transcriptome prior to NMP with renal blood flow at 2 hrs. A -** Comparison of enrichment scores for correlation pre and post 2 hrs NMP with urine output by GSEA against the Hallmarks database. Only pathways which are significant in at least 1 comparison have been plotted. Pathways which are altered between the conditions are labelled. **B** – Heatmap of all significant genes for correlation of transcriptome at 0hr with RBF at 2 hrs.

#### Supplemental Figure 3

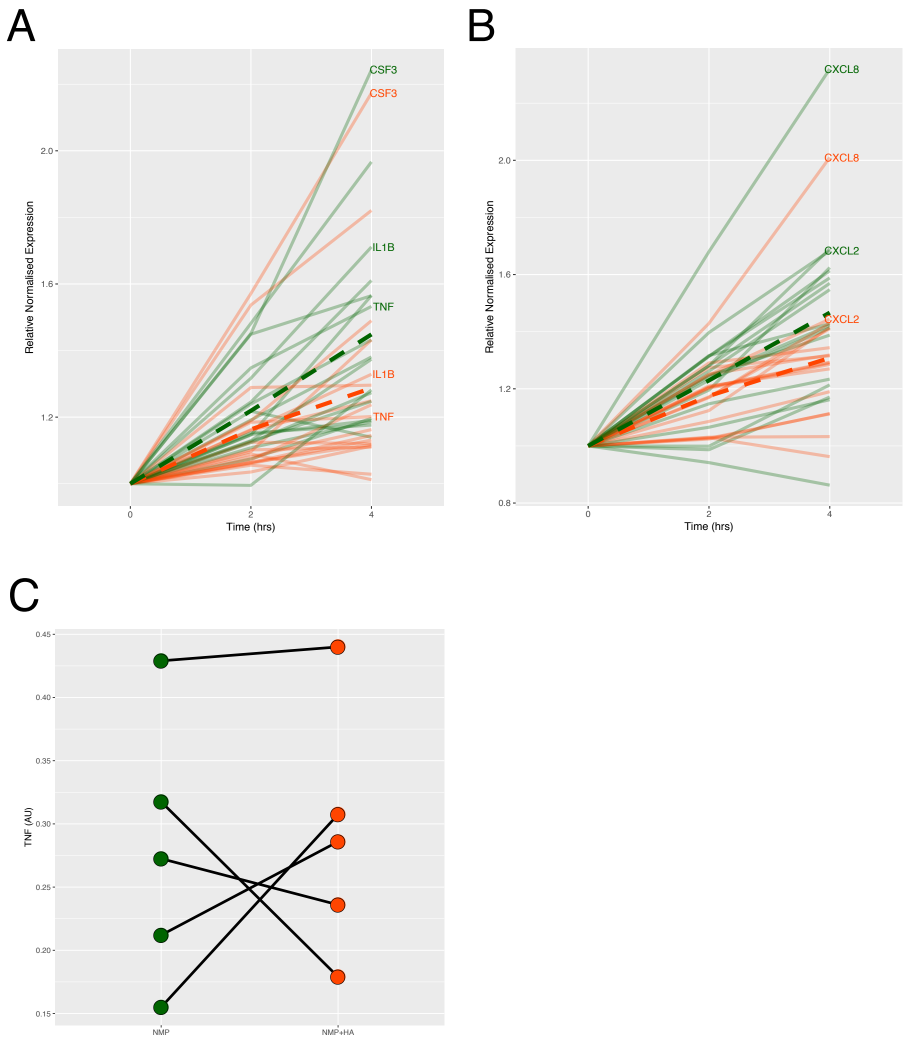


**Supplemental figure 3 – Effect of HA on cytokines and chemokines A** – Normalised gene expression profiles for all cytokines which were significantly altered by 4hrs post NMP. Selected cytokines annotated. Dashed line indicates LOESS regression line. Expression was normalised to respective time 0 for each group. Green line indicates NMP alone and orange with the addition of the haemoadsorber (HA). **B** – Normalised gene expression profiles for all chemokines which were significantly altered by 4hrs post NMP. Annotation as for A. **C** – Relative abundance of TNF protein in the tissue after 4 hrs NMP. The line indicates pairs of kidneys.
